# Diabetic conditions induce intolerance to accumulation of pathogenic mitochondrial DNAs

**DOI:** 10.1101/790956

**Authors:** Emi Ogasawara, Shun Katada, Takayuki Mito, Jun-Ichi Hayashi, Kazuto Nakada

**Affiliations:** Faculty of Life and Environmental Sciences, University of Tsukuba, 1-1-1 Tennoudai, Tsukuba, Ibaraki 305-8572, Japan; Graduate School of Life and Environmental Sciences, University of Tsukuba, 1-1-1 Tennoudai, Tsukuba, Ibaraki 305-8572, Japan; Life Science Center, Tsukuba Advanced Research Alliance, University of Tsukuba, 1-1-1 Tennoudai, Tsukuba, Ibaraki 305-8577, Japan

**Keywords:** Mitochondrial DNA, pathogenic mutation, Mitochondrial diseases, Diabetes, Model mice

## Abstract

Marked accumulation of mitochondrial DNA (mtDNA) with a particular pathogenic mutation is necessary for the mutant mtDNA to express its pathogenicity as mitochondrial respiration defects. However, the nuclear genome background, or the physiological status, or both, might also be important for the pathogenic regulation of mutant mtDNAs, because most mitochondrial function is controlled by polypeptides encoded in the nuclear genome. To test this, we generated diabetic mice carrying pathogenic mtDNA with a large-scale deletion (ΔmtDNA) that loses six tRNA genes and seven structural genes essential for mitochondrial respiration. Compared with non-diabetic mice carrying ΔmtDNA, diabetic mice carrying ΔmtDNA showed a decrease in mitochondrial biogenesis regulated by nuclear-encoded genes, and mitochondrial respiration defects and the resultant mitochondrial disease phenotypes were induced even in the case of low loads of ΔmtDNA. In addition, diabetic culture conditions intensified the pathogenicity of human mtDNA with an A3243G point mutation in the *tRNA*^*Lue (UUR)*^ gene. Our results indicated that the diabetic conditions are a modifier that exacerbates mitochondrial respiration defects due to mutant mtDNAs. The finding suggests the possibility that recovery from diabetic conditions might be an effective treatment strategy for some disorders involving both mutant mtDNAs and diabetic signs.

**Author Summary:** It has been reported that accumulation of pathogenic mutant mitochondrial DNA (mtDNA) and the resultant mitochondrial metabolic dysfunction are associated with a wide variety of disorders, such as mitochondrial diseases, diabetes, neuo-degenerative disorders, and cancers. Considering that most mitochondrial function is regulated by nuclear-genome-encoded polypeptides, it is very important to focus on cooperation between mutant mtDNA, nuclear genetic background, and vital conditions for understanding precise pathogeneses of mtDNA-mediated disorders. By using model cells and mice carrying pathogenic mtDNAs, we report here that diabetic conditions are a modifier for the pathogenic regulation of mutant mtDNAs. Because the onset and progression of diabetes are often associated with aging, our finding suggests that some age-associated disorders with mutant mtDNAs and diabetic complications might be induced partly by enhancement of the pathogenicity of mutant mtDNAs by diabetic conditions.

## Introduction

Mammalian mitochondria have their own genome, mitochondrial DNA (mtDNA), and most cells in the body contain multiple copies of mtDNA molecules (10^3^ to 10^4^ copies/cell). Mammalian mtDNA encodes 13 polypeptides that are essential subunits for electron transport complexes on the inner mitochondrial membrane, as well as 22 tRNAs and 2 rRNAs that are necessary for the translation of these 13 polypeptides. It has been well documented that pathogenicity of mutant mtDNAs is induced as mitochondrial respiration defects only when particular pathogenic mutant mtDNAs accumulate to substantial levels in affected tissues. This pronounced accumulation of pathogenic mtDNAs with large-scale deletions or particular point mutations leads to mitochondrial respiration defects that are manifested as a variety of mitochondrial diseases with clinical outcomes. We currently know that accumulation of mutant mtDNAs is not restricted to patients with mitochondrial diseases but extends to those with diabetes, neurodegenerative diseases, or cancers and further to aged subjects [1].

The diversity and onset of mtDNA-based disorders are considered to depend mainly on the magnitude and tissue distribution of mutant mtDNAs [1]. However, mitochondrial functional polypeptides—except for the 13 abovementioned ones that are derived from mtDNA—are encoded in the nuclear DNA. Therefore cooperation between mutant mtDNAs and the individual nuclear background or physiological status of the body might also be involved in the diversity and onset of mtDNA-based disorders. In fact, it has been reported that being homoplasmic for a pathogenic mutant mtDNA does not always induce serious clinical phenotypes in a mother and her children [2], even though the mother and her children carry the same pathogenic mtDNA because of maternal inheritance of mammalian mtDNA. At present, there is no experimental evidence as to whether the nuclear genome background and the body’s physiological status are less strongly the pathogenic regulation of mutant mtDNAs.

Patients with mitochondrial diseases who carry mutant mtDNAs frequently show diabetic phenotypes [3, 4]. Pathogenic ΔmtDNA and mtDNA with an A3243G point mutation in the *tRNA*^*Leu(UUR)*^ gene (3243mtDNA), which is associated with typical mitochondrial diseases [5, 6], have been found in families with maternally inherited diabetes and deafness (MIDD) [7, 8]. Moreover, abnormal mitochondrial metabolism, including abnormal mitochondrial respiration and fatty acid oxidation, has been observed in insulin-resistant states [9–11]. Therefore, many clinical studies have suggested that accumulation of pathogenic mutant mtDNAs and the resultant mitochondrial metabolic dysfunction are key contributors to diabetes (3), although it is accepted that diabetes is strongly associated with lifestyle and obesity [12, 13].

Mouse models that are heteroplasmic, with wild-type mtDNA and ΔmtDNA (See Fig *S1*), do not show a diabetic phenotype, but they have low blood glucose concentrations and high glucose sensitivities [14], even when they express mitochondrial diseases due to marked accumulation of ΔmtDNA throughout the body. Similarly, other mouse models with mitochondrial respiration defects specific to skeletal muscles or the liver do not have diabetic phenotypes [15, 16]. Moreover, a premature-aging model mouse carrying mtDNAs with multiple mutations due to disruption of the proofreading function of mtDNA polymerase does not show a diabetic phenotype [17]. Taken together, these results from model mouse studies suggest that mitochondrial dysfunction does not trigger the onset of diabetes. In addition, the loading proportion of 3243mtDNA increases along with age and the duration of the diabetic state [18], and, in diet-induced insulin-resistant mice, mitochondrial alterations do not precede the onset of insulin resistance and result from increased reactive oxygen species (ROS) production in the skeletal muscles [19]. These studies suggest that mtDNA mutation and mitochondrial dysfunction in diabetes might be consequences of the pathogenesis of diabetes. Therefore, whether mutant mtDNAs and mitochondrial dysfunction can cause diabetes remains unclear.

Here, we generated diabetic mice carrying ΔmtDNA to examine our hypotheses regarding the cooperation of mutant mtDNAs and individual nuclear backgrounds or physiological status, or both, in the pathogeneses of mitochondrial diseases and diabetes. We first hypothesized that the coexistence of mutant mtDNAs with the diabetic physiological state would result in pathogenicity manifesting as the exacerbation of diabetic phenotypes. We found instead that diabetic mice carrying ΔmtDNA escaped serious diabetic phenotypes, indicating that accumulation of ΔmtDNA did not lead to the exacerbation of diabetic phenotypes. We then hypothesized that the diabetic condition was responsible for the onset of mitochondrial diseases. We observed mitochondrial respiration defects and the onset of mitochondrial diseases in diabetic mice with low loads of ΔmtDNAs, indicating that the diabetic condition could, in fact, enhance the pathogenicity of ΔmtDNA. In addition, our *in vitro* experiments using human cybrids carrying 3243mtDNA showed that the pathogenicity of 3243mtDNA was enhanced by diabetic culture conditions. Our results therefore provide experimental evidence that the diabetic condition is a modifier for the pathogenic regulation of mutant mtDNAs.

## Results

### Accumulation of ΔmtDNA does not exacerbate diabetic phenotypes in mice with the db/db nuclear background

To elucidate whether mutant mtDNAs exerted pathogenicity manifesting as the exacerbation of diabetic phenotypes, we used diabetic db/db mice carrying 0% to 79% ΔmtDNA (db/db-mtΔ mice) in their tails (n = 27). “db/db” indicates a diabetic nuclear background derived from C57BLKS/J-+Lepr^*db*^/+Lepr^*db*^ mice and is a typical model for insulin-independent (type II) diabetes. db/db-mtΔ mice carrying 0% ΔmtDNA were used as a diabetic control.

We examined blood glucose concentrations and glucose sensitivities in the db/db-mtΔ mice. Lower blood glucose concentrations (Fig 1A) and higher glucose sensitivities (Fig 1B) were clearly observed in db/db-mtΔ mice with ≥49% ΔmtDNA, when compared with db/db mice carrying 0% ΔmtDNA. These results indicated that ΔmtDNA did not exacerbate diabetic signs: rather, mice with high levels of ΔmtDNA escaped the serious diabetic phenotypes caused by the db/db nuclear background.

**Fig 1.**
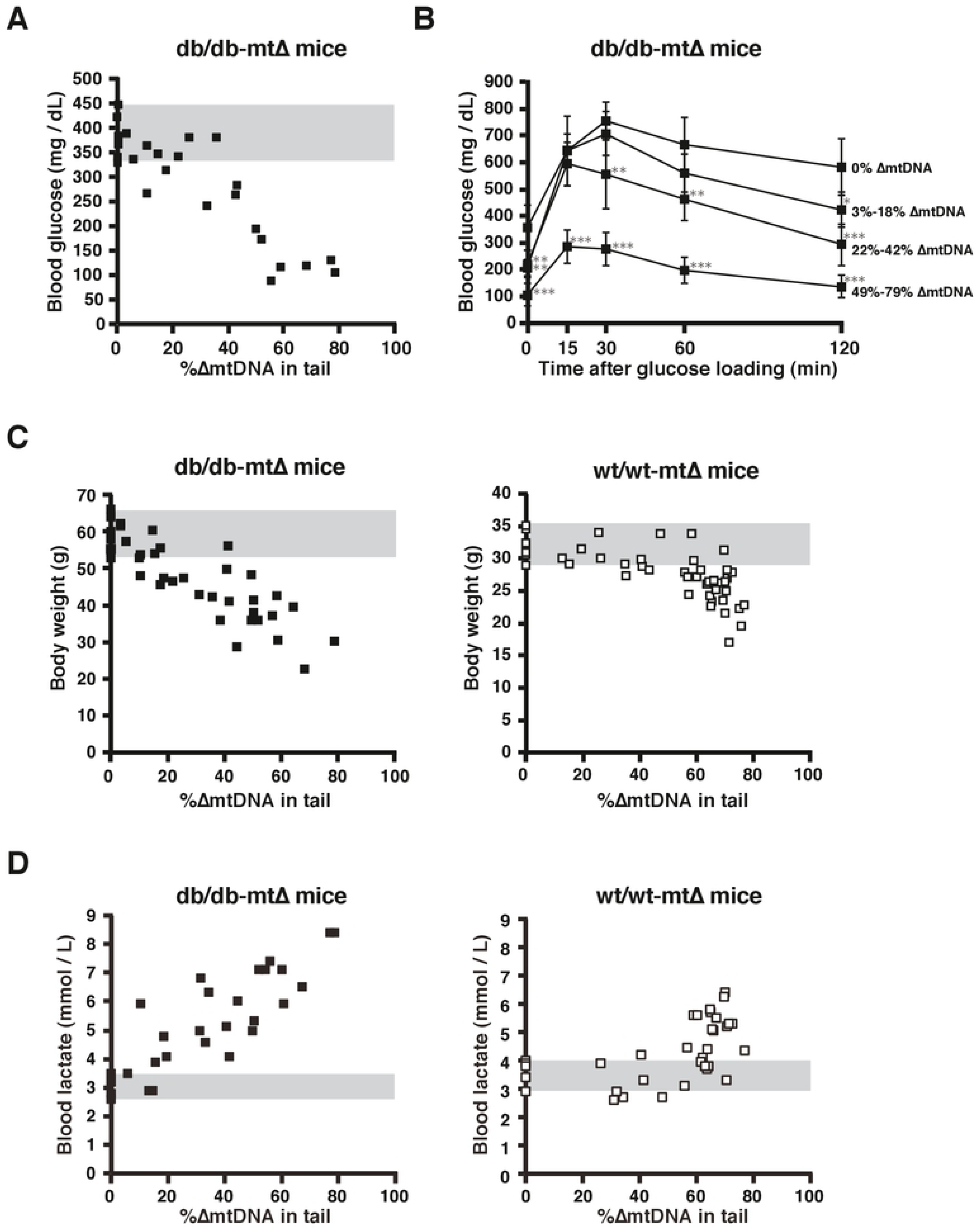
Phenotypic observations in db/db-mtΔ and wt/wt-mtΔ mice. (A) Non-fasting blood glucose concentrations in db/db-mtΔ mice. A decrease in non-fasting blood glucose concentration was clearly observed when the percentage of ΔmtDNA in the tail exceeded about 50%. (B) Glucose tolerance tests in db/db-mtΔ mice. Significant recovery from glucose intolerance was observed in db/db-mtΔ mice carrying 49% to 79% ΔmtDNA in their tails. (C) Comparison of body weights between db/db-mtΔ (left panel) and wt/wt-mtΔ mice (right panel). Compared with wt/wt-mtΔ mice, db/db-mtΔ mice showed a decrease in body weight, even with a low percentage of tail ΔmtDNA. (D) Comparison of blood lactate concentrations between db/db-mtΔ (left panel) and wt/wt-mtΔ mice (right panel). Unlike wt/wt-mtΔ mice, db/db-mtΔ mice showed onset of lactic acidosis, even with a low percentage of tail ΔmtDNA. Gray bands in (A), (C), and (D) indicate distributions of values in mice carrying 0% ΔmtDNA. All values are means ± 1 SD. *\*P* < 0.05; *\*\*P* < 0.01; *\*\*\*P* < 0.001. Statistical analysis was performed by one-way ANOVA with post hoc Dunnett’s testing for each time point.

### Diabetes enhances the pathogenicity of ΔmtDNA in mice

To elucidate whether the diabetic condition was responsible for the pathogeneses of mitochondrial diseases, we examined clinical signs of the onset of mitochondrial diseases in wt/wt-mtΔ mice carrying 0% to 77% ΔmtDNA (n = 38) and in db/db-mtΔ mice with 0% to 79% ΔmtDNA (n = 27); “wt/wt” indicates the wild-type nuclear background derived from C57BL/6 mice. Unlike wt/wt-mtΔ mice, db/db-mtΔ mice had low body weight (Wilk’s lambda = 0.3165, *P* < 0.0001) and lactic acidosis (Wilk’s lambda = 0.5990, *P* < 0.0001)—that is, clinical signs of the onset of mitochondrial diseases in wt/wt-mtΔ mice (20)—even when the ΔmtDNA load in db/db-mtΔ mice was lower than that in wt/wt-mtΔ mice (Fig 1C and 1D). We then performed histochemical analyses of the activities of electron transport complexes II (succinate dehydrogenase, SDH) and IV (cytochrome *c* oxidase, COX); cells showing mitochondrial respiration defects were visualized as negative for both nuclear DNA (nDNA)- and mtDNA-regulated COX activity and positive for nDNA-regulated SDH activity (*i.e.*, COX–/SDH+). Compared with those in wt/wt-mtΔ mice, the numbers of COX–/SDH+ cells in cardiac muscle tissues were clearly increased in db/db-mtΔ mice (Wilk’s lambda = 0.4295, *P* < 0.001), even when the ΔmtDNA load in the cardiac muscle tissues from db/db-mtΔ mice was almost equal to that in the tissues from wt/wt-mtΔ mice (Fig 2A and 2B). The same results were obtained from analyses of renal tissues (Wilk’s lambda = 0.7602, *P* <0.05) (Fig S2C and S2D). These observations suggested that mitochondrial respiration defects and the resultant mitochondrial diseases were frequently induced in db/db-mtΔ mice, even when these mice carried a low load of ΔmtDNA that could not induce mitochondrial respiration defects in wt/wt-mtΔ mice possessing a non-diabetic nuclear background.

**Fig 2.**
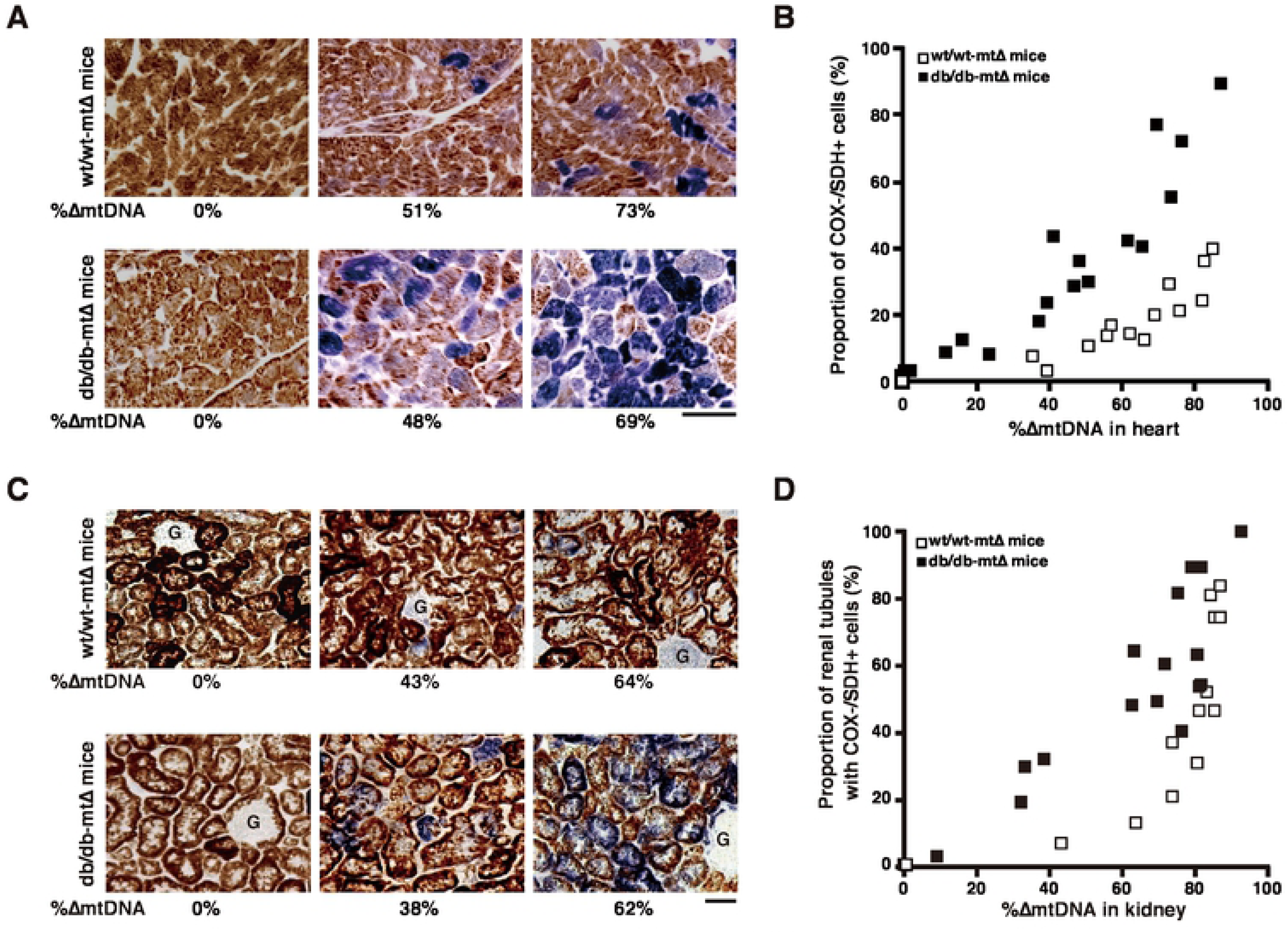
Comparison of mitochondrial respiration defects in db/db-mtΔ and wt/wt-mtΔ mice. Cells that were normal or deficient in mitochondrial respiration activity were visualized as brown (COX+/SDH+) or blue (COX–/SDH+), respectively. (A) Histochemical observations of mitochondrial respiratory enzyme activity in cardiac muscle tissues. The number of blue cardiomyocytes increased in db/db-mtΔ mice, even when the proportion of ΔmtDNA in cardiac muscle tissues from db/db-mtΔ mice was lower than that in tissues from wt/wt-mtΔ mice. Scale bar, 50 µm. (B) Relationship between ΔmtDNA load and proportion of cardiomyocytes showing mitochondrial respiration defects in cardiac muscle tissues. The proportion of cardiomyocytes with mitochondrial respiration defects in db/db-mtΔ mice was always higher than that in wt/wt-mtΔ mice. (C) Histochemical observations of mitochondrial respiratory enzyme activity in renal tissues. The number of blue cells increased in db/db-mtΔ mice, even when the proportion of ΔmtDNA in renal tissues from db/db-mtΔ mice was lower than that from wt/wt-mtΔ mice. Note that, because of a paucity of mitochondria, cells of glomeruli (G) were not stained sufficiently by the histochemical method used to show COX and SDH activities. Scale bar, 50 µm. (D) Relationship between ΔmtDNA load and proportion of renal tubules with cells showing mitochondrial respiration defects in renal tissues. Proportions of renal tubules with cells showing mitochondrial respiration defects were higher in db/db-mtΔ mice than in wt/wt-mtΔ mice. Scale bar, 50 µm.

Because db/db mice are typical models for type II diabetes, the results from experiments using db/db-mtΔ mice showed that the pathogenicity of ΔmtDNA was enhanced by the nuclear background or the physiological state of type II diabetes, or both. However, it was unclear whether a physiological state of type I diabetes could also enhance the pathogenicity of ΔmtDNA in mice. To clarify this point, we initially tried to generate wt/wt-mtΔ mice with streptozotocin (STZ)-induced type I diabetes, but wt/wt-mtΔ mice carrying more than 70% ΔmtDNA in their tails died by 7 days after STZ injection. This suggested that insulin secretion was necessary for the survival of wt/wt-mtΔ mice carrying a marked accumulation of ΔmtDNA. In the next experiment, we therefore used wt/wt-mtΔ mice carrying less than 70% ΔmtDNA and generated two kinds of mice: phosphate-buffered saline (PBS)-treated normal mice carrying 0% to 67% ΔmtDNA in their tails (PBS-treated wt/wt-mtΔ mice) (n = 15) and STZ-induced type I diabetic mice carrying 0% to 60% ΔmtDNA in their tails (STZ-treated wt/wt-mtΔ) (n = 15). The PBS-treated wt/wt-mtΔ mice were used as normal controls, and the STZ-treated wt/wt-mtΔ mice carrying 0% ΔmtDNA were used as type I diabetic controls. Because the blood glucose concentrations of STZ-treated wt/wt-mtΔ mice (509 ± 97.2 mg/dL) were significantly higher than those of PBS-treated wt/wt-mtΔ mice (143 ±15.0 mg/dL), we judged that the STZ-treated wt/wt-mtΔ mice expressed type I diabetes.

In the PBS-treated wt/wt mice, a tendency toward slight weight loss and lactic acidosis occurred when the loading proportion of ΔmtDNA in their tails reached approximately 60% (Fig S2A and S2B). Compared with PBS-treated wt/wt-mtΔ mice, STZ-treated wt/wt-mtΔ mice had significantly lower body weight (Wilk’s lambda = 0.591, *P* < 0.01; Fig S2A) and significantly greater blood lactate levels (Wilk’s lambda = 0.397, *P* < 0.001; Fig S2B). The numbers of COX–/SDH+ cells in the cardiac muscle and renal tissues were clearly greater in STZ-treated wt/wt-mtΔ mice (Wilk’s lambda = 0.698, *P* < 0.01 in cardiac muscle; Wilk’s lambda = 0.775, *P* < 0.05 in renal tissues) than in the PBS-treated wt/wt-mtΔ mice, even when the ΔmtDNA loads in these tissues from STZ-treated wt/wt-mtΔ mice were almost equal to those in the tissues from PBS-treated wt/wt-mtΔ mice (Fig S2C−F). From these results and those in the db/db-mtΔ mice, we concluded that the pathogenicity of ΔmtDNA was enhanced by not only type II, but also type I, diabetes in mice.

### Molecular regulation for induction of mitochondrial respiration defects due to even low loads of ΔmtDNA

Peroxisome proliferator-activated receptor gamma coactivator 1-alpha (PGC1α) is a positive regulator of mitochondrial biogenesis that controls mitochondrial transcription, translation, and replication [21]. PGC1α-responsible genes are coordinately downregulated in the state of insulin resistance and diabetes in humans [22, 23]. We therefore examined PGC1α-mediated mitochondrial biogenesis in the cardiac muscle tissues of wt/wt-mtΔ and db/db-mtΔ mice. Irrespective of the ΔmtDNA load, expression levels of PGC1α mRNA in the cardiac muscle tissues of db/db-mtΔ mice were significantly lower than those in wt/wt-mtΔ mice (Fig 3A). This suggested downregulation of mitochondrial transcription factor A (Tfam), which regulates mitochondrial transcription and replication downstream of PGC1α. In fact, the levels of Tfam mRNA in cardiac muscle tissues were significantly lower in db/db-mtΔ mice than in wt/wt-mtΔ mice (Fig 3A). We then examined the expression of mtDNA-encoded cytochrome *c* oxidase II (mt-Co2) mRNA and mtDNA copy numbers in the cardiac muscle tissues of wt/wt-mtΔ and db/db-mtΔ mice. mt-Co2 mRNA levels and mtDNA copy numbers in cardiac muscle tissues were significantly lower in db/db-mtΔ mice than in wt/wt-mtΔ mice (Fig 3A and 3B).

**Fig 3.**
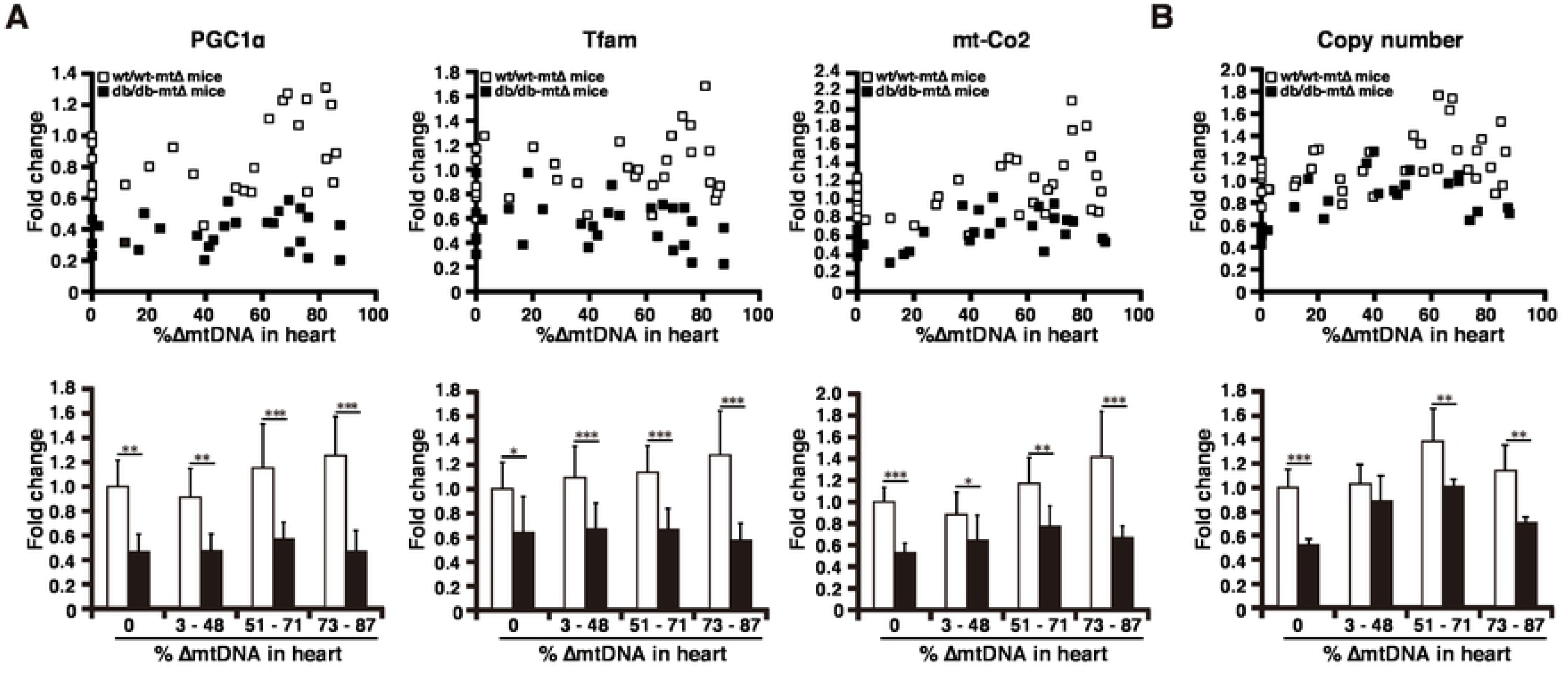
Changes in mitochondrial biogenesis in db/db-mtΔ mice and wt/wt-mtΔ mice. In (A) and (B), the upper graphs show plots of all values and the bottom ones show averaged values of four groups carrying 0%, 3% to 48%, 51% to 71%, and 73% to 87% ΔmtDNA in the cardiac muscle tissues. Statistical analysis was performed by using a *t*-test for comparison with wt/wt-mtΔ mice in each group. (A) Expression levels of PGC1α, Tfam, and mt-Co2 mRNAs in cardiac muscle tissues of wt/wt-mtΔ mice carrying 0% to 86% ΔmtDNA (open squares and bars) and db/db-mtΔ mice carrying 0% to 87% ΔmtDNA (solid squares and bars). Levels of PGC1α, Tfam, and mt-Co2 mRNAs in cardiac muscle tissues from db/db-mtΔ mice were significantly lower than in those from wt/wt-mtΔ mice. (B) Changes in mtDNA copy numbers in cardiac muscle tissues of wt/wt-mtΔ mice (open squares and bars) and db/db-mtΔ mice (solid squares and bars). Compared with those in wt/wt-mtΔ mice, mtDNA copy numbers in the cardiac muscle tissues from db/db-mtΔ mice were significantly lower, except for a case of 3% to 48% ΔmtDNA. In the bottom graphs, all values are means ± SD. *\*P* < 0.05; *\*\*P* < 0.01; *\*\*\*P* < 0.001.

These results indicating that PGC1α-mediated mitochondrial biogenesis was decreased in db/db-mtΔ mice are important for understanding why low loads of ΔmtDNA can induce mitochondrial respiration defects in db/db-mtΔ mice. The ΔmtDNA introduced into wt/wt-mtΔ and db/db-mtΔ mice loses six tRNA genes and seven structural genes. (See Fig S1) The main reason for the mitochondrial respiration defects due to marked accumulation of ΔmtDNA is insufficient mitochondrial translation for all 13 polypeptides encoded by mtDNA, owing to the shortage of six tRNAs containing the deleted region. Thus, a decrease in PGC1α-mediated mitochondrial biogenesis could exacerbate insufficient mitochondrial translation and the resultant mitochondrial respiration defects in db/db-mtΔ mice.

### Pathogenic regulation of human mtDNA with an A3243G point mutation in the tRNA^Lue(UUR)^ gene in diabetic culture conditions

As mentioned above, the pathogenicity of mouse ΔmtDNA was clearly enhanced by diabetes. However, whether the pathogenicity of human mutant mtDNA would also be altered by diabetic conditions was unclear. Because long-term culture in hyper-glucose medium is commonly used to create *in vitro* diabetic conditions [24, 25], we cultured two kinds of HeLa cybrid—namely a cybrid carrying only wild-type mtDNA (HeLa-mtWT) and a cybrid carrying 89% ± 1.3% mtDNA with an A3243G point mutation in the *tRNA*^*Lue(UUR)*^ gene (3243mtDNA) (HeLa-mtA3243G)—under normal and diabetic conditions. In cytochemical observations of COX and SDH activity (Fig 4A and 4B) and a biochemical analysis of COX activity (Fig 4C), HeLa-mtWT gave the same results in normal and diabetic culture conditions, suggesting that diabetic conditions did not affect mitochondrial respiration in these cybrids. Because the HeLa-mtA3243G cybrids used in this experiment possessed 89% ± 1.3% 3243mtDNA, they showed mitochondrial respiration defects even under normal culture conditions (Fig 4A–C). The numbers of HeLa-mtA3243G cells showing mitochondrial respiration defects (visualized as blue; COX–/SDH+) were significantly greater under diabetic culture conditions than in normal culture conditions (Fig 4A and 4B). The biochemical COX activity of HeLa-A3243G under diabetic culture conditions was significantly lower than that under normal conditions (Fig 4C). These data indicated that the pathogenicity of human mutant mtDNA, which is responsible for MELAS (mitochondrial myopathy, encephalopathy, lactic acidosis, and stroke-like syndrome) and MIDD [5–8], could be enhanced by diabetic culture conditions.

**Fig 4.**
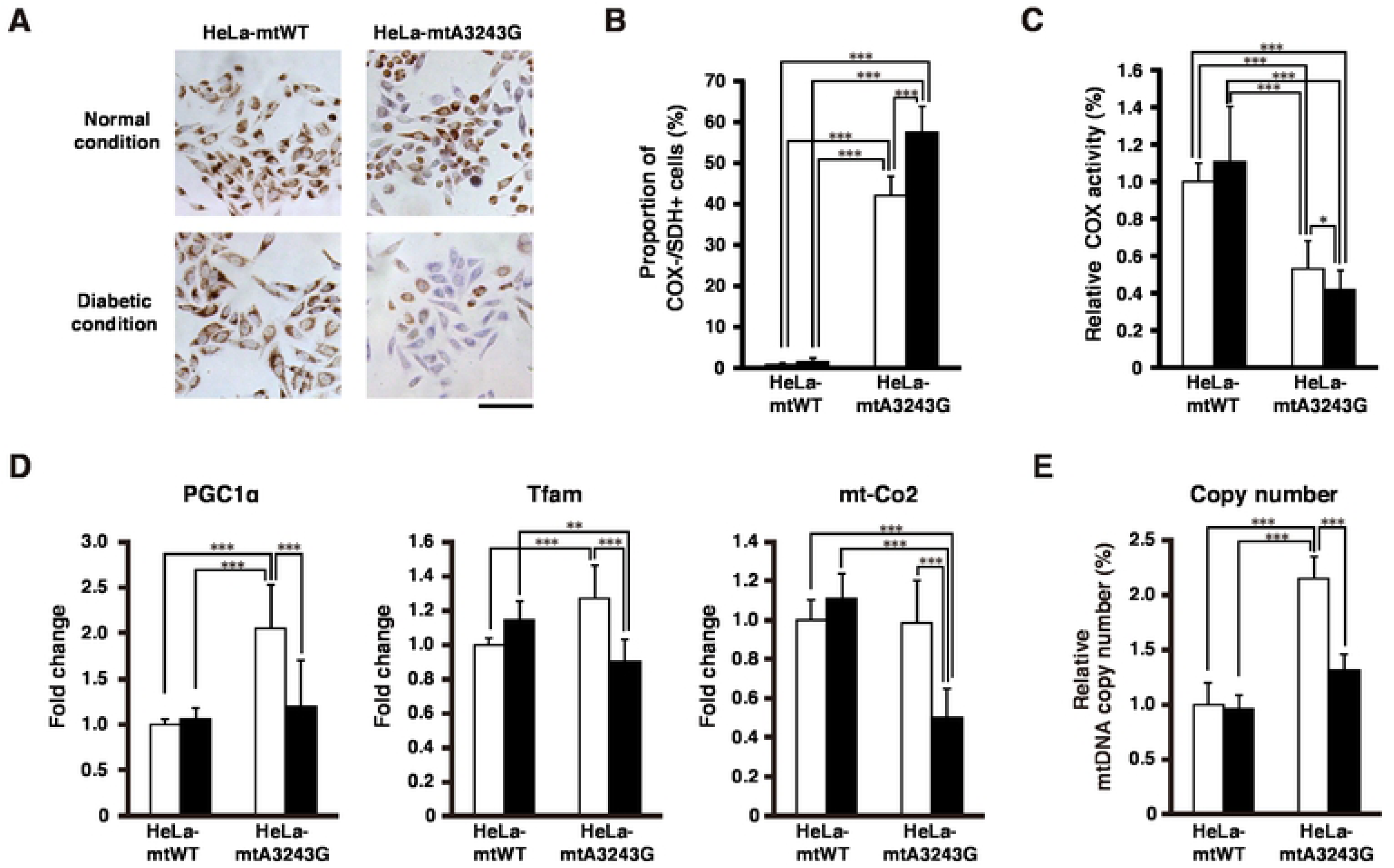
Characterization of human cybrids carrying mtDNA with the A3243G mutation in diabetic culture conditions. In this experiment, two human cybrids obtained by the repopulation of mtDNA-free HeLa cells with only wild-type mtDNA (HeLa-WT cybrids) or with both wild-type mtDNA and pathogenic A3243G mtDNA (HeLa-A3243G cybrids) were used. The HeLa-A3243G cybrids carried 89% A3243G mtDNA. The cybrids were cultured for 7 days with normal medium or high-glucose medium (representing diabetic culture conditions). In (B) to (E), open and solid bars indicate normal and diabetic culture conditions, respectively. (A) Cytochemical observations of COX and SDH activities. Cybrids that were normal or deficient in mitochondrial respiration activity were visualized as brown (COX+/SDH+) or blue (COX–/SDH+), respectively. The number of blue HeLa-A3243G cybrids increased, but that of HeLa-WT cybrids did not, when the cybrids were cultured in high-glucose medium. Scale bar, 50 µm. (B) Quantification of number of cybrids showing mitochondrial respiration defects. In comparison with culture in normal medium, the proportion of HeLa-A3243G cybrids visualized as blue (COX-/SDH+) increased significantly in culture with high-glucose medium. (C) Biochemical analysis for COX activity. The biochemical COX activity of HeLa-A3243G cybrids was half that of HeLa-WT cybrids in normal culture condition, because the HeLa-A3243G cybrids harbored large amounts of A3243G mtDNA (89% load). Culture in diabetic conditions further decreased the biochemical COX activity in HeLa-A3243G cybrids. (D) Levels of PGC1α, Tfam, and mt-Co2 mRNAs in HeLa-wt and HeLa-A3243G cybrids. HeLa-A3243G cybrids showed significantly lower levels of PGC1α, Tfam, and mt-Co2 mRNAs in diabetic culture conditions than in normal culture conditions. (E) Copy numbers of mtDNA in HeLa-wt and HeLa-A3243G cybrids. HeLa-A3243G cybrids in diabetic culture had significantly lower mtDNA copy numbers than in normal culture. All values are means ± 1 SD. *\*P* < 0.05, *\*\*P* < 0.01, *\*\*\*P* < 0.001. Statistical analysis was performed by using two-way ANOVA with post hoc Tukey’s HSD testing.

We next examined changes in the PGC1α-mediated mitochondrial biogenesis of HeLa-mtWT and HeLa-mtA3243G cultured under normal and diabetic culture conditions (Fig 4D and 4E). In the case of HeLa-mtWT, there were no significant differences in the levels of PGC1α, Tfam, and mtCo2 mRNAs, or in mtDNA copy numbers, between normal and diabetic culture conditions. In the comparison between HeLa-mtWT and HeLa-mtA3243G cultured in normal medium, significantly greater levels of PGC1α and Tfam mRNAs and mtDNA copy numbers were observed in HeLa-mtA3243G, suggesting that PGC1α-mediated mitochondrial biogenesis was accelerated to compensate for the mitochondrial respiration defects due to accumulation of 3243mtDNA, although there was no significant difference in the mt-Co2 mRNA levels between HeLa-mtWT and HeLa-mtA3243G under normal conditions. These HeLa-mtA3243G cybrids with accelerated PGC1α-mediated mitochondrial biogenesis, however, showed mitochondrial respiration defects in normal culture medium (Fig 4A–C). Importantly, the levels of PGC1α, Tfam, and mtCoII mRNAs and the mtDNA copy numbers of HeLa-mtA3243G were significantly lower in diabetic culture conditions than in normal culture conditions (Fig 4D and 4E). This indicated that diabetic culture conditions blocked the compensatory acceleration of PGC1α-mediated mitochondrial biogenesis in HeLa-mtA3243G. Because 3243mtDNA transcribes abnormal tRNA^Lue(UUR)^ [26], marked accumulation of 3243mtDNA induces a lack of normal tRNA^Lue(UUR)^, thus leading to insufficient mitochondrial translation and consequent mitochondrial respiration defects. Therefore, blockade of the compensatory acceleration of PGC1α-mediated mitochondrial biogenesis due to diabetic culture conditions could enhance the pathogenicity of human 3243mtDNA.

## Discussion

From the results of many clinical studies, it is well accepted that accumulation of pathogenic mutant mtDNAs, or mitochondrial metabolic dysfunction, or both, are responsible for the pathogenesis of diabetes [27, 28]. In addition, patients with mitochondrial diseases due to mutant DNAs frequently exhibit diabetic phenotypes [4]. In contrast, many model mouse studies have shown that pathogenic mtDNAs and mitochondrial dysfunction do not cause diabetes [14–17]. To gain an understanding of the multiple pathogeneses associated with mutant mtDNAs, we focused on the potential cooperation between mutant mtDNAs and the individual nuclear background or physiological status, or both, in the pathogeneses of diabetes and mitochondrial diseases. We first examined our hypothesis that pathogenic mtDNAs would have additional pathogenicity, thus exacerbating diabetic phenotypes, but not pathogenicity for the onset of diabetes. The phenotypes of db/db-mtΔ mice clearly indicated that accumulation of ΔmtDNA did not result in additional pathogenicity causing exacerbation of diabetic phenotypes. Therefore, our results, and those of previous study [14], suggested that marked accumulation of pathogenic mouse ΔmtDNA across the whole body was not associated with the onset and progression of diabetes. In contrast, it has been reported that mitochondrial dysfunction specific to pancreatic beta cells is responsible for diabetic phenotypes [29], and that ROS-generating mutant mtDNA induces a tendency toward age-related hyperglycemia in mice [30]. These findings, together with our results, suggest that the occurrence of diabetic phenotypes differs not only among the different types of mutant mtDNAs, but also among the tissue-specific distribution of mutant mtDNAs. Considering this point, it is possible that tissue-specific loading of ΔmtDNA could cause diabetes, although the distribution of ΔmtDNA over the whole body was not a trigger for the onset and progression of diabetes. To examine this possibility, we are now generating mice with tissue-specific distributions of ΔmtDNA.

Expression profiling studies have reported that PGC1α-responsible genes are coordinately downregulated in humans with insulin resistance and diabetes [22, 23]. To date, it has been considered that abnormalities in mitochondrial biogenesis, such as abnormalities in mtDNA replication, transcription, and translation, and the resultant mitochondrial dysfunction are a possible reason for the onset and progression of insulin resistance and diabetes. Consistent with the results of previous expression profiling studies [22, 23], we observed in the comparison between wt/wt-mtΔ mice and db/db-mtΔ mice carrying 0% mtDNA that the diabetic condition was associated with downregulation of PGC1α-mediated mitochondrial biogenesis (Fig 3A and 3B). Interestingly, we also observed that db/db-mtΔ mice carrying various proportion of ΔmtDNA showed further downregulation of PGC1α-mediated mitochondrial biogenesis and serious mitochondrial dysfunction (Fig 2 and Fig 3) but escaped the progression of diabetic phenotypes, even when they carried high loads of ΔmtDNA (Fig 1). These observations suggest that downregulation of PGC1α-mediated mitochondrial biogenesis in diabetes might not be behind the pathogenesis of diabetes, although because of biological differences we cannot extrapolate these results to humans.

Mitochondrial respiration defects due to the accumulation of ΔmtDNA can induce a compensatory increase in glycolysis and resultant lactic acidosis, because pyruvate is inadequately utilized at the level of reoxidation of reduced cofactors through the electron transport chain, and the glycolytic pathway is consequently accelerated to produce glycolytic ATP without mitochondrial respiration. Lactic acidosis in db/db-mtΔ mice, therefore, was indicative of the accelerated production of glycolytic ATP in these mice (Fig 1D). In contrast, exacerbation of diabetic phenotypes in db/db-mtΔ mice was not observed, even when these mice carried high loads of ΔmtDNA; instead, ΔmtDNA had the opposite effect of protecting these mice against serious diabetic phenotypes associated with their db/db nuclear genome background (Fig 1A and 1B). These results suggest that deconditioning of mitochondrial respiration and the resultant activation of glycolysis could counteract the pathogenesis of diabetes. In fact, imidodicarbonimidic diamide (metformin), a drug effective in the treatment of diabetes, can inhibit mitochondrial respiration [31].

We also hypothesized that diabetes would be associated with the pathogenesis of mitochondrial diseases. Our results provided direct experimental evidence that the diabetic condition can contribute to the onset of mitochondrial diseases, because the pathogenicity of ΔmtDNA was enhanced in mice with type I or II diabetes, and the onset of mitochondrial disease was induced even when the loading percentage of ΔmtDNA was lower than the pathogenic threshold for inducing mitochondrial respiration defects in non-diabetic mice. Furthermore, the pathogenicity of 3243mtDNA, which is a genetic cause of MELAS and MIDD [5–8], was enhanced under diabetic culture conditions. These findings showed that diabetes induces intolerance to the accumulation of pathogenic mutant mtDNAs and thus suggest that diabetic conditions are a modifier that exacerbates mitochondrial respiration defects due to mutant mtDNAs and the resultant onset and progression of mitochondrial diseases. Continuous downregulation of PGC1α-mediated mitochondrial biogenesis in diabetes was able to induce serious mitochondrial dysfunction in mice or cells carrying ΔmtDNA and 3243mtDNA (Fig 3 and Fig 4), but it was unclear how PGC1α-mediated mitochondrial biogenesis was downregulated by diabetic conditions.

Diabetic symptoms and signs are common in patients with mitochondrial diseases due to pathogenic mtDNAs [3, 4], and this provides potential evidence regarding which pathogenic mtDNAs cause diabetes. Our model mouse study, however, suggest a new explanation that the diabetic conditions would cause the onset of mitochondrial diseases even in the case of low loads of mutant mtDNAs, rather than mutant mtDNAs causing diabetes, and that this probably leads to a high frequency of diabetes in a population with mitochondrial diseases. Of course, clinical support for this explanation requires further investigation.

In our *in vitro* experiment using human cybrids carrying 3243mtDNA, the pathogenicity of the 3243mtDNA was clearly enhanced by diabetic culture conditions (Fig 4A–4C). In a cohort study of patients with MELAS caused by the accumulation of 3243mtDNA, the frequency of diabetic symptoms was higher in older patients than in younger ones [32]. The implication is that phenotypes of MELAS in some older patients might be caused by their diabetes. In age-associated disorders, such as neurodegenerative diseases and cancer, as well as in aging, the accumulation of pathogenic mutant mtDNAs or mitochondrial abnormalities, or both, is frequently observed [8]. Because the onset of type II diabetes is often associated with aging [3], some age-associated phenotypes and disorders might be caused by the enhanced pathogenicity of mutant mtDNAs in diabetes. These findings provide the possibility that recovery from diabetes could be an effective treatment strategy, not only for some mitochondrial diseases, but also for some age-associated disorders with both accumulation of mutant mtDNAs and diabetic symptoms, although biological and clinical evidence for this treatment strategy remains to be accumulated.

## Materials and methods

### Mice

All animal experiments were performed in compliance with the University of Tsukuba’s institutional guidelines for the care and use of laboratory animals. db/db-mtΔ mice were obtained by the mating of male db/+ mice with female mito-miceΔ, because mammalian mtDNA is inherited maternally [33]. Assays of non-fasting blood glucose concentration, glucose tolerance, body weight, and blood lactate concentration were performed at about 8 months after birth. Tissue samples collected from the mice were used to estimate ΔmtDNA loadings and for histochemical analysis. wt/wt-mtΔ mice (n = 30) were also used for experiments using STZ-induced diabetic models.

### Generation of STZ-induced diabetic models

To generate STZ-induced diabetic models, wt/wt-mtΔ mice carrying 0% to 67% ΔmtDNA in their tails (n = 30) (10 to 12 weeks old) were used. Seventeen mice were injected with STZ solution (85 mg/kg body weight) (Sigma) via the tail vein for 2 consecutive days. The remaining 13 mice were injected with PBS solution and used as controls. Mice with blood glucose levels over 350 mg/dL at 2 weeks after STZ injection were used as models of STZ-induced diabetes. Changes in body weight and blood lactate concentration in these mice were examined at about 20 to 22 weeks after birth (10 weeks after STZ or PBS injection). Tissue samples were then collected and used for experiments.

### Cells and culture conditions

HeLa cybrids carrying only wt-mtDNA (HeLa-wt cybrids) or both wild-type mtDNA and pathogenic mtDNA with an A3243G mutation in tRNA^Leu(UUR)^ were used. HeLa-wt and HeLa-A3243G cybrids were cultured for 7 days in RPMI1640 (Nissui) containing 110 mM glucose with 0.1 mg/mL pyruvate, 50 μg/mL uridine, and 10% bovine serum albumin (BSA) as experimental diabetic conditions; cultures with high glucose levels are used to reproduce diabetic conditions in *in vitro* experiments [24, 25]. In addition, cybrids were cultured for 7 days in RPMI1640 (Nissui) containing 11 mM glucose with 0.1 mg/mL pyruvate, 50 μg/mL uridine, and 10% BSA as normal conditions. Culture media were changed every other day.

### Measurement of blood glucose and lactate concentrations

Blood glucose and lactate concentrations were measured with a glucometer (DEXTER-Z, Bayer) and lactate sensor (ARKRAY), respectively, in accordance with the manufacturer’s protocol. Blood samples collected from the tail vein were used for these measurements.

### Glucose tolerance tests

For glucose tolerance tests, mice were fasted overnight, and then a glucose solution (1.5 g/kg body weight) was orally loaded. Blood samples were collected from the tail vein, and glucose concentrations were measured with a glucometer (G Checker with Cyclic GB Sensor, Gunze).

### Estimation of ΔmtDNA load

Proportions of wild-type mtDNA and ΔmtDNA were determined by using a real-time PCR technique, as described previously [34]. Total DNA samples of tails, hearts, and kidneys were used in this experiment.

### Histochemical and cytochemical procedures

Histochemical and cytochemical analyses of the activities of nuclear-genome-encoded SDH (complex II in the mitochondrial respiratory chain) and mtDNA-encoded COX (complex IV in the mitochondrial respiratory chain) were performed as described previously [35, 36]. Cardiac muscle tissues carrying 0% to 85% ΔmtDNA from wt/wt-mtΔ mice or 0% to 87% ΔmtDNA from db/db-mtΔ mice, and renal tissues carrying 0% to 87% ΔmtDNA from wt/wt-mtΔ mice or 0% to 92% ΔmtDNA from db/db-mtΔ mice, were used. Cryosections (thickness, 10 μm) of these tissues were stained for COX activity and then stained for SDH activity. Two cultured cell lines, HeLa-wt and HeLa-A3243G cybrids, were used as additional samples for double staining for COX and SDH activities.

In this method, the activities of nuclear-genome-encoded SDH and mtDNA-encoded COX are visualized as blue and brown, respectively. In the case of mutant mtDNA-based mitochondrial respiration defects, COX activity decreases, whereas SDH activity compensatorily increases. Thus, cells showing mitochondrial respiration defects (COX–/SDH+ cells) are detected as blue. To estimate the cell population showing mitochondrial respiration defects, approximately 6.5-mm^2^ fields in each stained section were randomly selected under a microscope (model DMRE, Leica Microsystems) and the proportions of cardiomyocytes and renal tubules stained blue were calculated. Similarly, more than 7000 HeLa cybrids stained for COX and SDH activities were counted, and the proportion of HeLa cybrids showing mitochondrial respiration defects was estimated.

### Measurement of mtDNA copy number

Total DNAs containing both nDNA and mtDNA were isolated from cardiac muscle and renal tissues. mtDNA copy number was assayed by using a real-time PCR technique with Power SYBR Green (Applied Biosystems). Nuclear gene glyceraldehyde-3-phosphate dehydrogenase (GAPDH) was also assayed as an internal control, using the following specific primers: 5’CCCACGATCAACTGAAGCAGCAA3’ and 5’ACCGTTTGTTTGTTGAAATATT3’ for mouse mtDNA; 5’AACGACCCCTTCATTGAC3’ and 5’TCCACGACATACTCAGCAC3’ for mouse GAPDH; 5’TACATTACTGCCAGCCACCA3’ and 5’GTGGCTTTGGAGTTGCAGTT3’ for human mtDNA; and 5’TAGAGGGGTGATGTGGGGAG3’ and 5’AGTGATGGCATGGACTGTGG3’ for human GAPDH.

### Analysis of gene expression

Total RNAs were isolated from cardiac muscle and renal tissues with TRIzol reagent (Life Technologies), and reverse transcription was performed with a High Capacity cDNA Reverse Transcription Kit (Applied Biosystems) in accordance with the manufacturers’ instructions. Gene expression levels were assayed by using a Taqman gene expression assay with mouse PGC1α (Mm 01208835_m1) and mouse Tfam (Mm 00447485_m1) in real-time PCR analysis. The mRNA of β-actin as an endogenous control, mtCo2, human PGC1α, and human Tfam were assayed by using a Power SYBR Green (Applied Biosystems) with the following specific primers: 5’-GGTCATCACTATTGGCAACGAG-3’ and 5’-GTCAGCAATGCCTGGGTACA-3’ for mouse β-actin; 5’-GCACAAGAAGTTGAAACCA-3’ and 5’-CGGATTGGAAGTTCTATTGG-3’ for mouse mtCo2; 5’-GGAGACGTGACCACTGACAATGA-3’ and 5’-TGTTGGCTGGTGCCAGTAAGAG-3’ for human PGC1α; 5’-CCGAGGTGGTTTTCATCTGT-3’ and 5’-GCATCTGGGTTCTGAGCTTT-3’ for human Tfam; and 5’-TTCATGATCACGCCCTCATA-3’ and 5’-TAAAGGATGCGTAGGGATGG-3’ for human mtCo2.

### Statistical analysis

Data from the glucose tolerance tests at each time point were analyzed by using Dunnett’s test, and the values from db/db-mtΔ mice populations carrying various percentages of ΔmtDNA were compared with those from db/db-mtΔ mice carrying 0% ΔmtDNA. Data on gene expression and mtDNA copy numbers from the *in vivo* experiments using mice were analyzed by using Student’s *t*-test, and values from db/db-mtΔ mice populations were compared with those from wt/wt-mtΔ ones. To evaluate whether data on body weight, blood lactate, and cell population with mitochondrial respiration defects differed significantly between wt/wt-mtΔ mice carrying 0% to 77% ΔmtDNA and db/db-mtΔ mice carrying 0% to 79% ΔmtDNA in their tails, a step-wise forward discriminant function analysis was performed. Data from the *in vitro* experiments using human cybrids were analyzed by using Tukey’s HSD test. In these statistical analyses, JMP software (SAS Institute, Cary, NC) was used, and values with *P* < 0.05 were considered to differ significantly.

## Acknowledgments

This work was supported by Grants-in-Aids for Scientific Research (A) (No. 23240058) and (B) (No. 16H04678) from the Ministry of Education, Culture, Sports, Science, and Technology of Japan (MEXT) to K.N. and for Scientific Research (A) (No. 16H02463) from MEXT to J.-I.H. This work was also supported by the World Premier International Research Center Initiative of MEXT (to K.N. and J.-I.H.). This work was also supported by AMED-CREST (18gm1110006 to K.N.) from the Japan Agency for Medical Research and Development.

## Supporting information

**S1 Fig. Gene maps of mouse mtDNA and ΔmtDNA.**

Mouse mtDNA encodes 37 genes, consisting of 13 polypeptides, 22 tRNAs, and 2 rRNAs (left panel). The ΔmtDNA loses seven structural and six tRNA genes owing to a large-scale deletion (right panel). The arc in ΔmtDNA indicates the deleted region, which is expanded from the tRNA^*Lys*^ to *ND5* genes. The wt/wt-mtΔ and db/db-mtΔ mice that we used in this experiment harbored both wild-type mtDNA and ΔmtDNA. Capital letters in the two maps indicate the 22 tRNA genes.

**S2 Fig. Phenotypic observations of wt/wt-mtΔ mice with STZ-induced diabetes.**

(A) Comparisons of body weight between PBS- (open symbols) and STZ-injected (filled symbols) wt/wt-mtΔ mice. (B) Comparisons of blood lactate concentration between PBS-(open symbols) and STZ-injected (solid symbols) wt/wt-mtΔ mice. Unlike PBS-injected wt/wt-mtΔ mice, STZ-injected ones showed onset of lactic acidosis even with a low load of ΔmtDNA. (C) Histochemical observations of COX and SDH activities in cardiac muscle tissues. Scale bar, 50 µm. (D) Relationship between ΔmtDNA load and proportion of cardiomyocytes showing mitochondrial respiration defects (cardiomyocytes stained blue). (E) Histochemical observations of mitochondrial respiratory enzyme activities in renal tissues. (F) Relationship between ΔmtDNA load and proportion of renal tubules with cells showing mitochondrial respiration defects (renal tubules stained blue). Scale bar, 50 µm. Gray bands in (A) and (B) indicate distributions of values in mice carrying 0% ΔmtDNA.

